# Q-SHINE: a versatile sensor for glutamine measurement via ligand-induced dimerization

**DOI:** 10.1101/2022.06.28.497868

**Authors:** Moon-Hyeong Seo, Yun Lim, Ji Yul Kim, Youn Hee Jung, Jae Hoon Lee, Min Seok Baek, Je Hyeong Jung, Ho-Youn Kim, Wookbin Lee, Keunwan Park

## Abstract

Studies on glutamine (Gln) metabolism have illuminated the vital role of Gln in cellular functions and its potential as a biomarker for disease detection. Despite the increasing interest in Gln metabolism, in-depth evaluations are challenging owing to limitations of conventional Gln-measuring methods. Thus, we developed a ligand-induced dimerization-based sensor for Gln, termed Q-SHINE, by splitting a glutamine binding protein into two separate domains. Q-SHINE enables highly accurate and convenient measurement of Gln concentration in bio-fluid samples, and the detection range is optimal for physiological Gln levels. Genetically encoded Q-SHINE sensors could also visualize intracellular Gln levels and quantify cytoplasmic and mitochondrial Gln change in living cells, which enabled detection of various cell responses to extracellular Gln supplement.

## 1. Introduction

Glutamine (Gln) is a versatile amino acid that participates in the synthesis of nucleic acids, proteins, and anabolic processes to produce carbon and nitrogen. The optimal concentration of Gln is crucial in maintaining cellular integrity and function [1,2]. Previous studies have reported alterations in Gln levels in several diseases including cancer [2,3], diabetes[4,5], and neurodegeneration [6–8]. In particular, the consumption rate of Gln in cancer cells is significantly higher than that in normal cells, which indicates that Gln metabolism may be a key target for cancer treatment [2,3,9,10]. Therefore, supplementation or deprivation of Gln has been suggested to treat these diseases [2,7,11–14]. Previous studies have provided crucial insights into the significance of Gln levels in cellular functions and the potential application of Gln as a diagnostic biomarker of the relevant pathologies.

Measurement of Gln levels in physiological samples has previously been performed using analytic equipment (high-performance liquid chromatography [15–17] and amino acid analyzer [18]) or an enzyme-based colorimetric Gln assay [11,12]. However, the requirement of intricate preparation steps, such as cell lysis, deprotonation, and derivatization, hinders expanded use of these methods [19]. Furthermore, the accuracy of Gln assay kits are affected by underlying glutamate levels or other endogenous compounds interfering the enzymatic reactions. Alternatively, Gln concentration can be measured with spatiotemporal resolution in a genetically encoded manner via fluorescence-coupled sensors (based on FRET or circularly permuted fluorescent proteins) using a glutamine-binding protein (QBP) [20–23]. However, low dynamic range and frequent experimental errors limit their widespread adoption in the Gln metabolism research and diagnostic applications.

Here, we report a novel glutamine-sensing protein, Q-SHINE (Gln sensor via Split HINg-Emotion binding protein), constructed by splitting a QBP via cleavage of two lobes. After divided, we confirmed the two independent domains of QBP reconstitute upon Gln binding. This ligand-induced hetero-dimerization (LID) system was subsequently coupled to a desired reporter, such as mCherry or NanoLuc, to transform the binding event into a detectable signal. Our LID-based Gln sensors, Q-SHINE_Red for fluorescence, and Q-SHINE_Luc for bioluminescence, provide a simplified protocol and higher specificity than conventional Gln assay kits, facilitating reagentless point-of-care diagnoses for diverse diseases related to Gln metabolism. Genetically encodable Q-SHINE sensors effectively visualize intracellular Gln levels, and further enable comparative quantification of cytoplasmic and mitochondrial Gln changes in response to the environment of cancer and non-cancer cell lines.

## 2. Materials and Methods

### 2.1. Split design of PBPs

The domain movement of proteins was analyzed using DynDom [35]. In the case of QBP, two heterogeneous QBP conformers were compared (PDB ID: 1GGG and 1WDN). Results showed that QBP consists of two motional domains and exhibits an open-closed bending motion (rotation angle: 55.7 degree, and closure ratio: 99.8%). The residues situated in the hinge-bending region were 87–89 and 181–184 in the QBP (1WDN). Splitting the QBP at the two connecting strands generated two separate domains, where one domain (QBP_Lg) was fragmented into two segments. The discontinuous fragments were redesigned by modeling a new linker sequence, which connected the cleaved ends to generate a single polypeptide chain. Linker modeling was performed using RosettaRemodel with default options [36]; loop closure trials were attempted 10,000 times for loop lengths ranging from two to five. The designed loops were then filtered using LoopAnalyzerMover in RosettaScript [37]. The Rosetta total score, which summarizes loop quality, was used to select the final loop sequence that stabilized the fused domain. A shorter loop length was also prioritized to minimize flexibility. Structures of the designed domains were predicted using AlphaFold Colab [38].

### 2.2. Gene cloning for bacterial, mammalian, and plant cell expressions

Genes for the sensor components (QBP, mCherry, NanoBiT, and other PBPs with nucleotide sequences complementary to pET21a and Novagen at both ends) were synthesized by IDT gBlocks^®^ Gene Fragments (Integrated DNA Technologies, Iowa, USA) and Gene Fragments (Twist Bioscience, South San Franscisco, USA). For bacterial expression, the pET21a plasmid was linearized through restriction with the NdeI and XhoI enzymes. The synthesized gene fragments were then cloned into the linear pET21a vector using the In-Fusion^®^ HD Cloning Kit (Takara Bio, Kusatsu, Japan). For mammalian expression, the pcDNA3.1 (+) vector was linearized using PCR amplification. The split QBP and mCherry constructs were amplified, assembled, and cloned using the In-Fusion HD Cloning Kit. For plant cell expression, the whole expression cassette from the pcDNA plasmid was cloned into the pTALCOMT plasmid for Agrobacterium-mediated transformation [39].

### 2.3. Protein expression and purification

Plasmids containing the sensor constructs were transformed into *E. coli* BL21 (DE3) cells. Protein expression was induced with 0.5 mM isopropyl b-D-1-thiogalactopyranoside at 18 °C. Proteins were purified from the soluble cell lysate using His60 Ni Superflow Resin (Takara Bio) and concentration was measured using NanoDrop. For analytical gel filtration chromatography, the purified proteins from a Ni column were loaded onto a Superdex 75 Increase column equilibrated with 20 mM Tris-HCl (pH 8.0, 150 mM NaCl). Molecular mass standards included conalbumin (75 kDa), ovalbumin (44 kDa), carbonic anhydrase (29 kDa), and ribonuclease (13.7 kDa).

### 2.4. *In vitro* BiFC

Next, 5 μM of both sensor domains were mixed with different concentrations of Gln or other amino acids in the Nunc Microwell 96-well Microplate (Thermo fisher, MA, US), and the mixture was incubated in the dark for 1–5 h at 22 °C. Emission spectra of mCherry were read in 1 nm increments from 600 to 620 nm using an Infinite M1000 (Tecan, Männedorf, Switzerland) under 580 nm excitation.

### 2.5. Mammalian cell culture

Human embryonic kidney 293T (HEK293T) was purchased from the American Type Culture Collection (ATCC). The cell line was tested for mycoplasma contamination and used only within 30 passages. HEK293T was maintained in Dulbecco’s Modified Eagle Medium (DMEM; Gibco, Massachusetts, USA) supplemented with 10% (v/v) fetal bovine serum (FBS; Gibco), 100 U/mL penicillin and 100 μg/mL streptomycin (P/S; Cytiva) at 37 °C in a humidified 5% CO_2_ incubator. HeLa cells (a gift from Dr. Lee, KIST, Korea) were maintained in DMEM (Gibco) supplemented with 10% FBS (Gibco), and 1% PS at 37 °C.

### 2.6. Bioluminescence measurement

Next, 5 μM of two domains of Q-SHINE_Luc with 0–5 mM Gln for 10 min at 22 °C in the dark. We also added 25 μL of the Nano-Glo Live Cell reagent included in the Nano-Glo Live Cell Assay System (Promega) to each well and gently mixed for 10 s. Luminescence was measured for a single time point using the GloMax Navigator System (Promega) and an integration time of 1 s. For intracellular Gln quantification, cells were seeded at 1 × 10^4^ cells/well in white/clear flat-bottom 96-well plates (Corning) and pCDNA plasmids harboruing Q-SHINE_BL or mit-Q-SHINE_BL were transfected with 50 ng/well within 24 hours using Lipofectamine 3000 reagent (Invitrogen) according to the manufacturer’s instructions. For glutamine starvation, cell culture medium was changed with glutamine-free DMEM (Gibco) with 10% FBS before transfection and Opti-MEM medium (Gibco) was replaced with MEM (Welgene). Glutamine added to final concentration at 0–2 mM after 16 h glutamine starvation. Time-dependent bioluminescence was measured after glutamine addition using Nano-Glo^®^ Dual-Luciferase^®^ reporter assay system (Promega). Cell culture plates were equilibrated at 22 °C. Next, 80 μL/well of ONE-Glo^™^ EX Reagent was added into 80 μL/well cell culture medium and mixed on an orbital shaker (300 rpm) for 5 min at 22 °C. Then, firefly luciferase activity was measured by a GloMax Navigator Microplate Luminometer (Promega) with 1 s integration time. Subsequently, 80 μL NanoDLR™ Stop & Glo^®^ Reagent was added to each well, mixed for 10 min on an orbital shaker (600 rpm). NanoLuc activity also was measured by a GloMax^®^ Navigator Microplate Luminometer (Promega) with 1 s integration time. Data were analyzed using Prism 9.0 (GraphPad Software).

### 2.7. Measurement of Gln concentration in serum samples

Female C57BL/6 mice, aged 6–12 weeks, were purchased from Orient-Bio (Seongnam, Korea). The mice were cared for in accordance with institutional guidelines for experimental animals. All experimental protocols were approved by the Animal Care and Use Committee of KIST (Approval No. KIST-2021-09-104). To collect serum samples, blood was taken from the abdominal caval vein and centrifuged at 300 *×g* for 10 min at 4 °C after anesthesia was administered and an abdominal incision was created. At least 200 μL serum was collected and stored at −80 °C. For Gln measurement, 20 μL of serum samples from each mouse and different concentrations of Gln solution were dissolved in distilled water. The samples were then mixed with 5 μM of two domains of Q-SHINE_Red, followed by incubation in the dark and at 22 °C for 5 h. The standard curve is fitted by nonlinear curve fitting method using the Hill1 function in Origin 2020 (OriginLab Corporation, Northampton, USA), and the concentration of Gln in serum samples was calculated using the ‘Find X from Y’ function based on the fitted graph. For comparison, Gln concentration in mouse serum was simultaneously determined using the EnzyChrom Glutamine Assay Kit (EGLN-100, BioAssay Systems, Hayward, CA, USA) according to the manufacturer’s protocol.

### 2.8. Monitoring Gln changes in mammalian cells

The mammalian expression vector was transiently transfected with Lipofectamine 3000 (Invitrogen, Thermo Fisher Scientific, Massachusetts, USA). For the intracellular bimolecular fluorescence complementation (BiFC) experiments, HEK293T cells were seeded into 12-well cell-culture treated plates (Thermo Fisher Scientific) and grown at 37 °C in 5% CO_2_ overnight in DMEM supplemented with 10% FBS and 1% P/S to 4–50% confluency. After the medium was changed to DMEM (no Gln) supplemented with 10% FBS and 1% P/S to deprive the cells of endogenous Gln, cells were transfected with plasmids containing Q-SHINE_FL or Q-SHINE_BL proteins using Lipofectamine 3000. After 16 h of transfection, 0–2 mM of Gln were added to the media for induction of Gln-dependent BiFC. Live cells were imaged with the EVOS FL Cell Imaging System (Thermo Fisher Scientific) after the addition of Gln. To obtain separate images of GFP and mCherry, two light cubes (470 nm excitation, 525 nm emission; 530 nm excitation, and 593 nm emission) were used. Images from both channels were overlaid to visualize overall fluorescence. Fluorescence intensities of mCherry and GFP were quantified using Image J software (National Institutes of Health). The images were segmented into each color in grayscale and the intensity of each fluorescence was measured on a median scale after the ROI settings were established through threshold application.

### 2.9. Transient expression and imaging of fluorescent proteins in tobacco leaves

*Agrobacterium-mediated* transient expression analysis was performed as described by Lee et al [40]. *Agrobacterium tumefaciens* LBA4404 cells carrying different sensor constructs were adjusted to OD600 0.4 in resuspension solution (10 mM MgCl2, 10 mM MES-KOH (pH 5.6), 100 μM acetosyringone) and infiltrated into leaves of 4 to 5-week-old *Nicotiana tabacum* cv. Samsun. Plants were grown and incubated at 24/21 °C (day/night temperatures), for a 16 h photoperiod, with 500 μmol/m^2^/s light intensity and at 80 % relative humidity. At 5 days post-infiltration, fluorescence intensities from the leaves were determined using an LSM5 Zeiss confocal laser scanning microscope with the ZEN image analysis software (Carl Zeiss, Jena, Germany). The excitation/emission spectra were 488/493–598 nm for GFP and 594/599–650 nm for mCherry.

### 2.10. Statistical analysis

Data from at least three independent experiments were expressed as the mean ± standard deviation (SD). Statistical significance was determined using a one-way analysis of variance followed by Tukey’s multiple comparisons test using GraphPad Prism 9 (GraphPad Software).

## 3. Results and Discussion

### 3.1. Split design of a Gln-binding protein

QBP, a type II periplasmic binding protein (PBP), consists of two domains connected by a hinge wherein a ligand binds to the region between two lobes. Ligand binding induces the closing and twisting motions of the QBP that bring the lobes closer by hydrogen bonds between the two domains and the ligand. Based on this structural feature, we converted *E.coli* QBP into Gln-recognizing domains (Q-SHINE) that dimerize upon ligand binding (**Figure 1**). By comparing the open and closed structures (PDB: 1GGG and 1WDN), the domain movement of QBP through the hinge-bending motion was analyzed to estimate the cleavage sites that were supposed to cause the least structural damage to the resulting split domains, and to ensure minimal effect on ligand binding. Residues 87–89 and 181–184 (amino acid numbering in PDB 1WDN) were identified as hinge-bending regions and two domains were separated by the deletion of Lys87-Ser88 and Ala182 (**Figure 2A**). Cutting two connecting strands of the QBP results in noncontinuous fragment for a large domain (Lg; residues 1–86 and 183–226) and a continuous fragment for a small domain (Sm; residues 89–181); thus, a linker was required for the stable expression of the large domain. Of the three tested loops, a designed linker (TGNG) that closed the gap between Tyr86 and Gln183 in QBP Lg displayed superior solubility when expressed in *E.coli* (data not shown). AlphaFold showed the predicted structures of the two divided domains, in which was insignificantly different from the global fold of the native QBP (**Figure 2B**).

**Figure 1.**
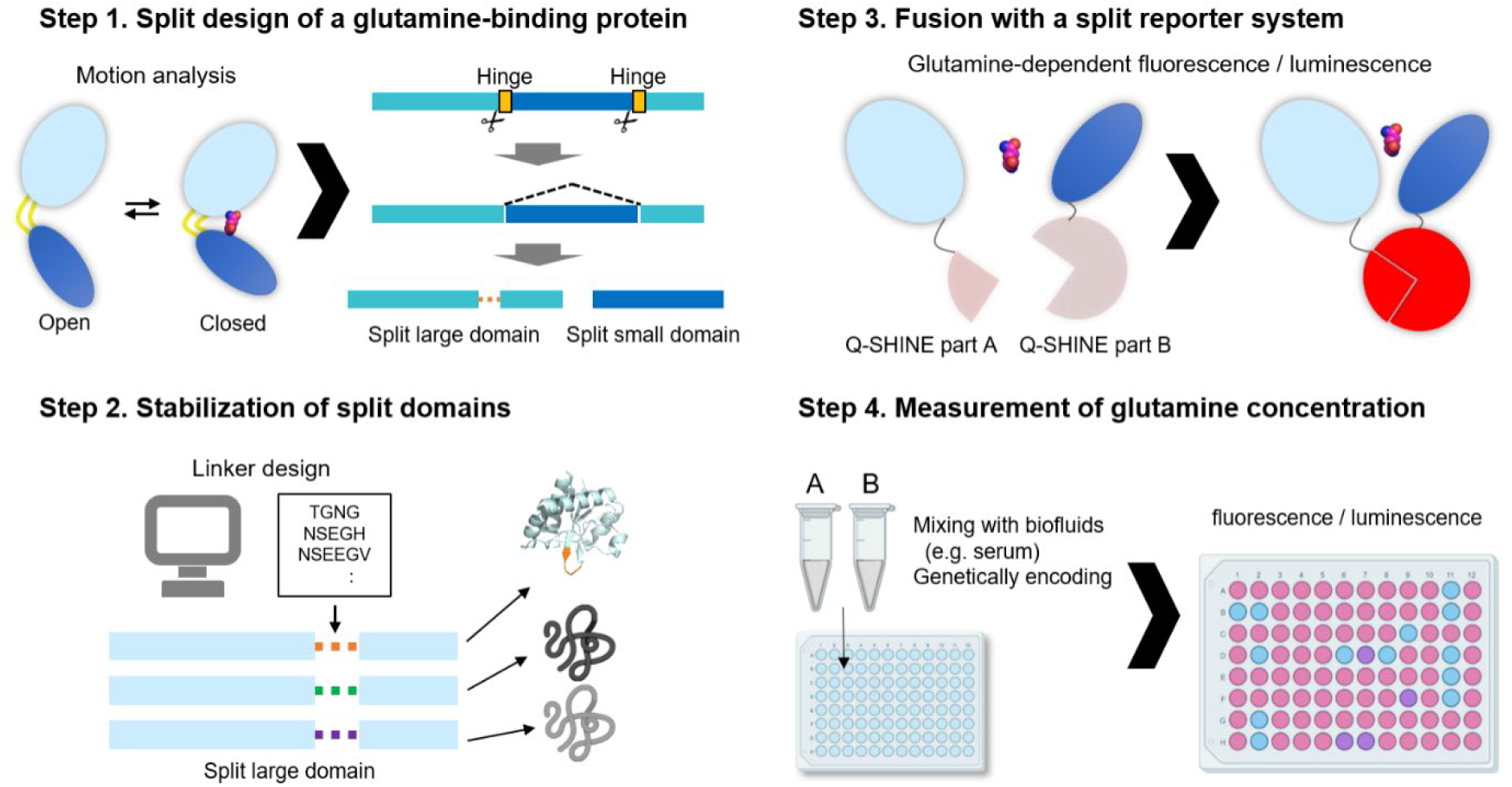
Schematic of Q-SHINE design and Gln measurement. In step 1, based on the open and closed structure of glutamine-binding protein (QBP), hinge residues are selected by analysis of domain movement. In step 2, for the stable expression and proper function, suitable linker sequence was designed for the noncontinuous fragment. In step 3, the separate domains are fused with split reporter domains, such as split fluorescent protein and luciferase. In step 4, two components of the final sensor are mixed with samples or genetically expressed in cells. Resulting signals (fluorescence or bioluminescence) represent Gln level.

**Figure 2.**
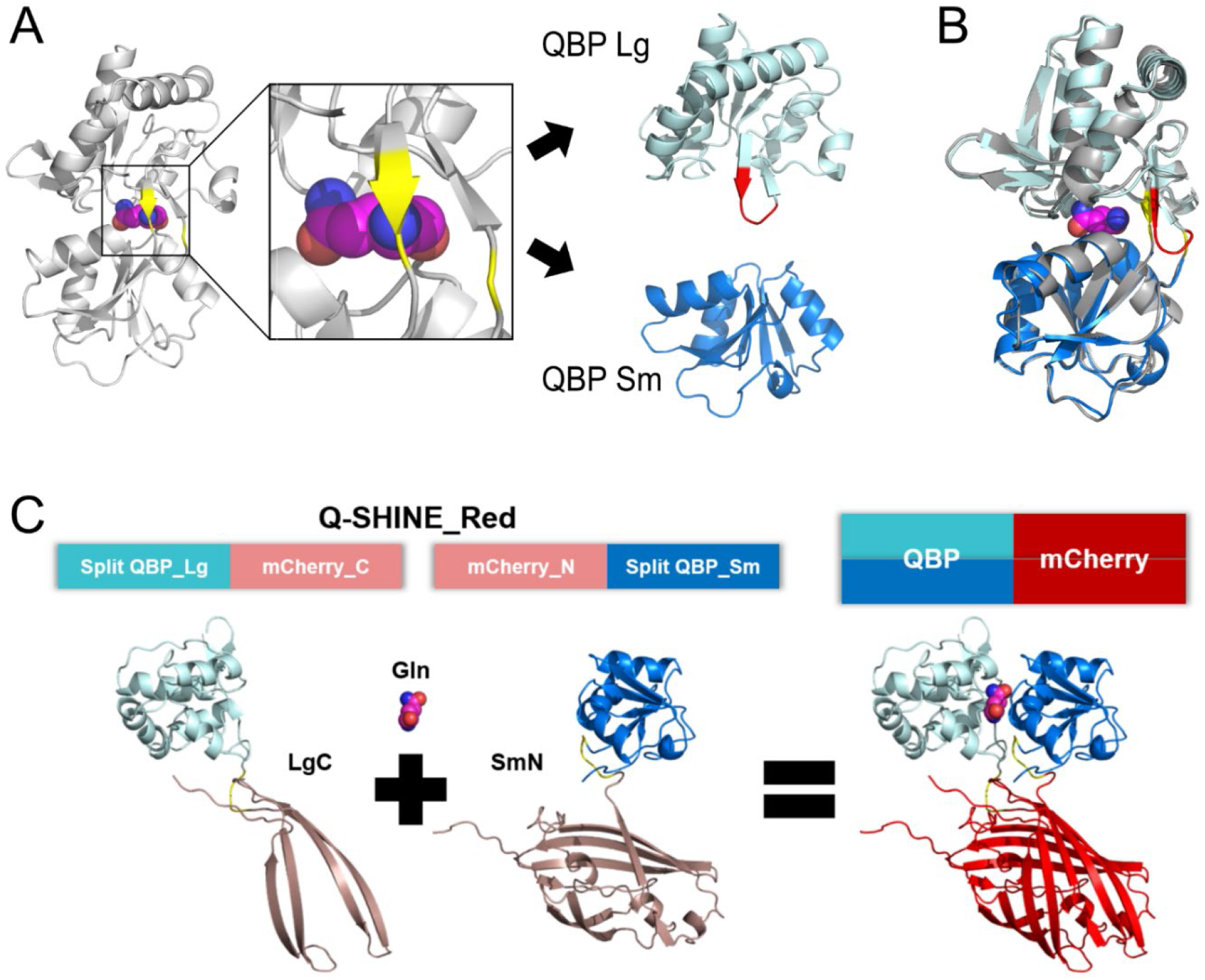
Design of split QBP and Q-SHINE_Red. (A) Based on the hinge-bending motion of *E. coli* QBP (PDB 1WDN), hinge residues (yellow) were selected for cleavage. One noncontinuous fragment was connected by the design of a new linker (red), resulting in two separate domains of QBP: QBP_Lg (light blue) and QBP_Sm (blue). (B) Structures of split domains were predicted by AlphaFold. When aligned with the holoprotein, two split domains showed the same global fold as in native QBP (RMSD 0.516 Å for QBP Lg, 0.418 Å QBP Sm). (C) Each split QBP domain was recombined with a split mCherry fragment for the construction of the complete Q-SHINE_Red system: Split QBP_Lg – mCherry_C (LgC) and mCherry_N – Split QBP_Sm (SmN). Gln-induced heterodimerization of QBP gives rise to red fluorescence by complementation of split mCherry fragments.

Each split QBP fragment, Lg and Sm, was subsequently recombined with split mCherry fragments (MN159 & MC160),[24] which serve as reporter elements, resulting in the final fusion construct, Q-SHINE_Red consisting of Split QBP_Lg-mCherry_C (LgC) and mCherry_N – Split QBP_Sm (SmN) (**Figure 2C**). Gly-Ser linkers were used to link the split QBP and mCherry fragments. Two domains of Q-SHINE_Red were analyzed first using gel filtration chromatography. Both domains (LgC and SmN) moved faster than their expected size (**Figure S1**), which might be because the nonnative structural configuration of the recombinant fragments, which is elongated rather than spherical (**Figure 2C**) since the two independent fragments are connected by a flexible linker. Interestingly, LgC was isolated as two separate peaks (LgC_1 and LgC_2) while SmN was eluted as a single peak. However, when either LgC_1 and LgC_2 was mixed with SmN in the presence of Gln, the signal-to-noise ratio (S/N; mCherry fluorescence from the well with the ligand to the without ligand) of each pair displayed insignificant differences. The complementation rate (the increase of fluorescence) was faster when LgC_2 was used (data not shown). Although the underlying molecular mechanism determining the difference of complementation efficiency remains elusive, LgC_2 was solely used in the subsequent experiments. Because split mCherry can associate without being brought into close proximity by connected domains, the association rate of two domains without Gln was examined by comparing Q-SHINE_Red and Q-SHINE_Red_ΔSm. Fluorescence of Q-SHINE_Red_ΔSm is thought to be dependent on the self-assembly of split mCherry domains merely because of QBP Sm deletion. Interestingly, Q-SHINE_Red exhibited a significantly slower increase of fluorescence in the absence of Gln by the attenuated self-assembly than that of Q-SHINE_Red_ΔSm (**Figure S2**). Fusion of the split QBP domain may hinder the self-complementation of split mCherry fragments, which is beneficial for a higher S/N ratio in measuring Gln concentration. Taken together, bimolecular fluorescence complementation (BiFC) of the split mCherry domains of Q-SHINE_Red should provide a qualitative and quantitative readout of the Gln-induced dimerization of split QBP domains, thereby enabling the measurement of Gln concentration.

Next, we examined whether the split design could be adopted in other Type II PBPs with structural similarity *(Campylobacter jejuni* cysteine-binding protein (CBP) [25] and *E. coli* histidine-binding protein (HBP) [26]). However, a split CBP with an identical design strategy showed limited dynamic range of Cys-dependent fluorescence and no His-dependent signal from split HBPs (**Figure S3).**Given that the reported binding affinity of HBP is higher than that of QBP and CBP (30, 300, and 100 nM, respectively), the imbalance in the binding mode of His against two domains of HBP might be accountable for the poor ligand-induced dimerization (**Table S2**). Although Q-SHINE uncovered the potential of a PBP family for a novel ligand-responsive split system, the elaborate choice of a PBP scaffold and subsequent design for split domains should be critical to the successful generation of a novel PBP-based sensor.

### 3.2. Gln measurement using Q-SHINE_Red

To assess the ability of Q-SHINE_Red to detect Gln in solution, we combined equivalent concentrations (5 μM) of Q-SHINE_Red proteins (LgC and SmN) with varying concentrations of Gln. After incubating for a few hours, Gln-induced QBP dimerization was recognized by red fluorescence in accordance with the proximity-induced complementation of mCherry (**Figure 3A**). While the signal contrast was maximized during the 3–6 h incubation period, the sensitivity increased with a longer incubation time. In terms of analytical sensitivity, Q-SHINE_Red could detect Gln concentrations as low as 1 μM. The emission peak was observed at 610 nm, which is the reported value for the native mCherry protein (**Figure 3B**). The ligand selectivity of Q-SHINE was further tested using structurally similar amino acids: glutamate, asparagine and D-Gln. As shown in **Figure 3C**, Q-SHINE exhibited a highly specific reactivity to Gln, clearly distinguishing it from other amino acids. Importantly, Q-SHINE_Red showed notable sensitivity in the range of hundreds of μM to mM concentrations, demonstrating its suitability for measuring physiological Gln levels which typically fall in the 200–1400 μM range [12,19,22,27].

**Figure 3.**
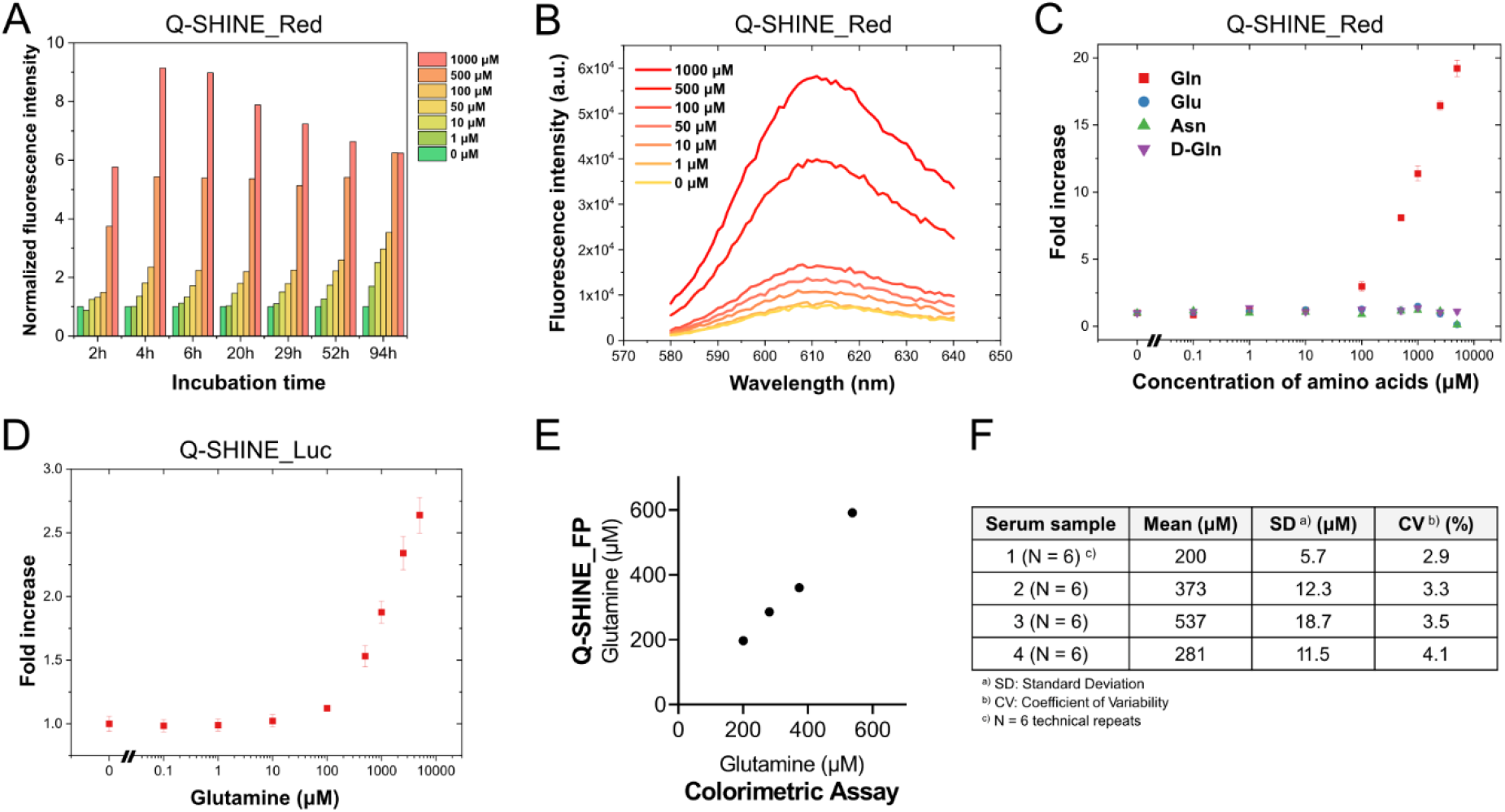
Gln-measurement with Q-SHINE_Red and Q-SHINE_Luc. (A) Gln-dependent BiFC signal was measured with the marked concentration of Gln 2–94 h of incubation. Each signal intensity was normalized to the well without Gln. (B) The representative emission spectra changes by different Gln concentration. (C) Ligand selectivity of Q-SHINE_Red was examined with other amino acids with similar structure. (D) Bioluminescence of Q-SHINE_Luc increased depending on the Gln concentration. The x-axis represents the corresponding Gln concentration in the reaction mixture. Data were normalized to the signal from the wells no amino acid added. Error bars represent the s.d. in triplicate experiments. (E) Comparison of Q-SHINE_Red against a commercially available Gln assay kit in measuring Gln concentration of mouse serum. Pearson’s correlation coefficient, R^2^ = 0.9938, *P* = 0.0062. (F) Intra-assay variability of Q-SHINE_Red for measuring Gln concentration in mouse serum.

We sought to examine the performance of Q-SHINE by exchanging the reporting element with a split luciferase. Unlike mCherry, NanoBiT is a reversible complementation system based on enzymatic reactions that enable real-time kinetic analysis of live cells [28]. The split QBP and NanoBiT fragments were recombined in a similar manner resulting in Q-SHINE_Luc (**Table S2**). Similar to the results of Q-SHINE_Red, an increase in the bioluminescence was observed in proportion to Gln concentration (**Figure 3D**). The sensitivity of the NanoBiT-based system was commensurate with that of the Q-SHINE_Red because the sensing elements are essentially identical in both systems. The Q-SHINE_Red showed a higher fold change in signals (i.e., high signal-to-noise ratio) owing to cumulative fluorescence by irreversibly complemented mCherry, while NanoBiT gave an immediate response in less than 10 min. This suggests that other split reporter systems [29–31] such as β-lactamase, kinase, protease, biotin ligase, or Cas9, could be readily combined with Q-SHINE for various applications.

We then investigated the performance of Q-SHINE_Red in quantifying Gln in a biological sample. The each Q-SHINE domain was combined with 20 μL each four different mouse sera as well as standard Gln solutions from 0 to 2 mM. Fluorescence intensities from the wells were read by excitation at 587 nm and emission at 610 nm 5 h after incubation. The Gln concentration of mouse sera was determined based on the standard curve obtained by plotting the values from standard Gln solutions (**Figure S3**). Q-SHINE_Red provided data comparable to a conventional and commercially available Gln colorimetric assay kit (**Figure 3E**). Furthermore, intra-assay analysis verified the high repeatability and accuracy of Q-SHINE_Red for the serum samples (**Figure 3F**). Cells, tissues, or clinical samples with high glutamate or protein contents often result in high background signals in conventional colorimetric Gln assays, which require additional pretreatment or background control. Endogenous compounds may also interfere the enzymatic reactions. Given the limited methodologies, Gln is one of the lesser-highlighted biomarkers despite the growing number of reports demonstrating its relevance in multiple diseases, including cancer [2,11,27], diabetes [4,5], and neurodegeneration [6–8]. The simple composition of Q-SHINE_Red, containing only two protein domains, enables straightforward measurement without any pretreatment process or expensive equipment. Q-SHINE_Red will therefore produce highly reliable and accurate data in biological and biomedical research that utilizes conventional Gln assay kits.

### 3.3. Monitoring Gln changes in living cells

We investigated whether genetically encoded Q-SHINE could monitor intracellular Gln change. For the equimolar expression of Q-SHINE fragments, a self-cleaving T2A peptide was inserted between the two elements. GFP was also attached as an internal control to normalize mCherry signal, resulting in Q-SHINE_FL (**Figure 4A**), which was expressed in Gln-deprived HEK293T cells. It has been shown that the intracellular amino acid concentrations are largely dependent on the amounts of amino acids in medium [32]. When different concentrations of Gln were added to the medium, red fluorescence was detected from cells fluorescing GFP as a control (**Figure 4B & Figure S4**). As expected, Q-SHINE-FL displayed red-to-green signal in proportion to the Gln concentration and the incubation time. (**Figure 4C**). We also tested in tobacco leaves to examine whether the biosensor could detect Gln in a different cellular environment, and Q-SHINE-FL successfully visualized Gln in the plant cells (**Figure S5**).

**Figure 4.**
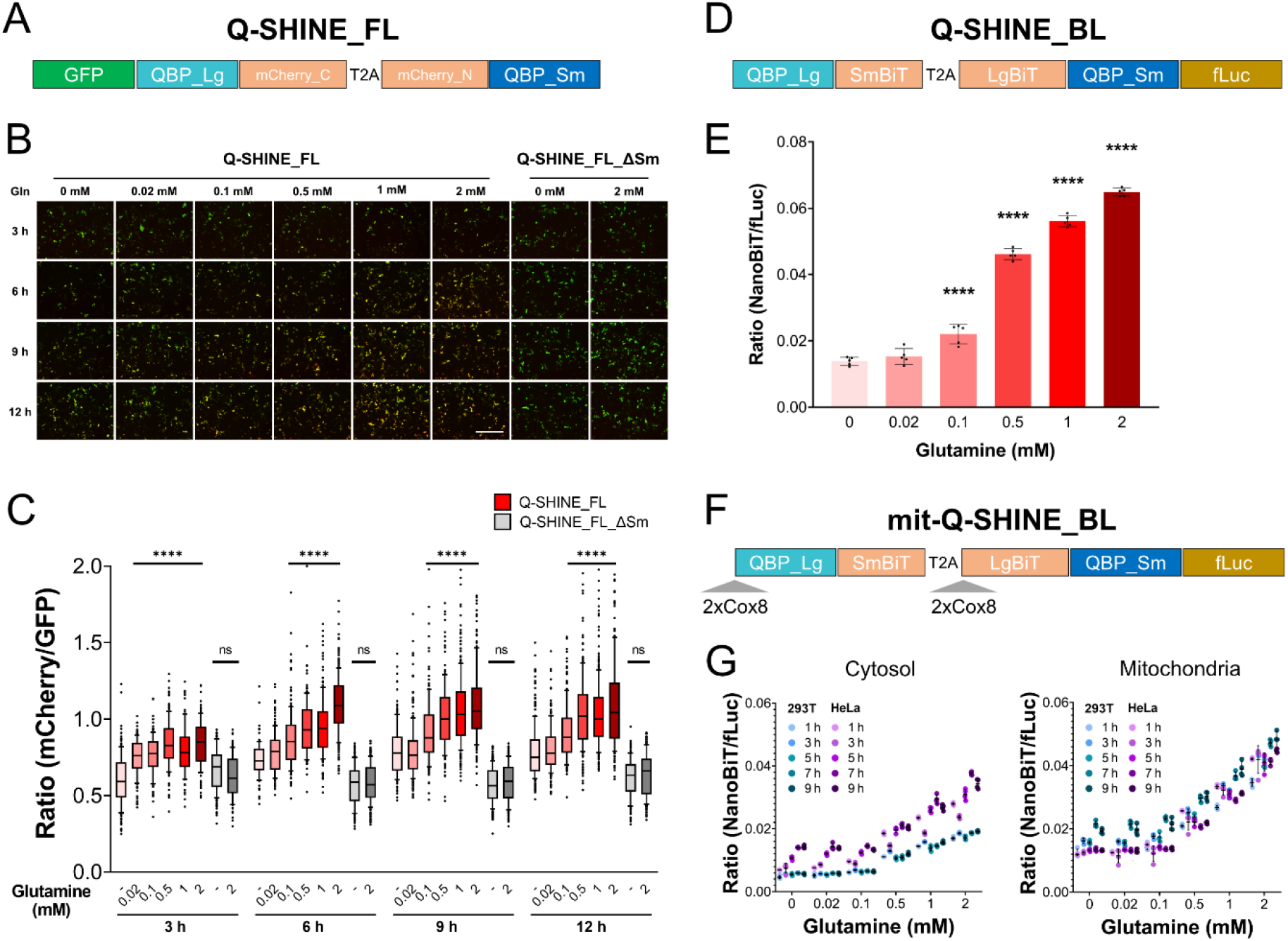
Monitoring Gln changes in living cells. (A) Schematic of the Q-SHINE_FL for the monitoring intracellular Gln level. (B) Fluorescence microscopy of HEK293T cells transfected with Q-SHINE-FL with increasing Gln concentrations. Overlay images from green and red channels are shown. Scale bars, 400 μm. (C) Image-based quantification of mCherry/GFP fluorescence intensity ratio by Q-SHINE_FL at different Gln concentrations. Gln- and time-dependent increase in the fluorescence ratio by Q-SHINE-FL was clearly detected while the ratio of Q-SHINE_FL_ΔSm remained unchanged. The number of ROIs ranges from 89 to 319. The levels of significance are indicated as follows: ****P< 0.0001; n.s., not significant. (D) Schematic of the Q-SHINE_BL for the quantitative analysis of intracellular Gln level. (E) Q-SHINE_BL exhibited highly consistent increase of NanoBiT/fLuc ratio depending on the Gln concentrations. (mean±s.d). (F) mit-Q-SHINE_BL was constructed by addition of 2xCox8 sequences at the N-terminal of final cleaved fragments of Q-SHINE_BL. (G) Bioluminescence ratio in HEK293T and HeLa was compared using Q-SHINE_BL and mit-Q-SHINE_BL by supplementing different Gln concentrations to Gln-starved cells. Error bars represent the s.d. in quadruplicate experiments.

For the quantitative analysis of intracellular Gln levels, we modified Q-SHINE_Luc into a genetically encoded form of Q-SHINE_FL, resulting in Q-SHINE_BL (**Figure 4D**). Gln-dependent bioluminescence ratio of NanoBiT to firefly luciferase (fLuc) from the live cells was observed as expected (**Figure 4E**). We then expressed the biosensor in mitochondrial matrix by adding two tandem copies of the mitochondrial targeting signal from cytochrome c oxidase subunit VIII to the N-terminus of both Q-SHINE_BL fragments [32] (**Figure 4F**). We compared mit-Q-SHINE_BL and Q-SHINE_BL signals in both HEK293T and HeLa cell lines to explore the potential of our biosensor to specifically monitor and discriminate Gln levels in both cytosol and mitochondria. Indeed, homeostasis of Gln metabolism maintained by transport of Gln across plasma or mitochondria membrane, cytoplasmic glutamine synthesis, and mitochondrial glutaminolysis under various physiological conditions is vital for normal function and proliferation of cells [1]. Bioluminescence ratio was measured with the live cells after adding Gln to the medium. After different concentrations of Gln were added, time-dependent change of Gln levels either in cytosol or mitochondria was compared in both cells. As a result, Gln transport into the cytosol and/or de novo Gln synthesis in cytosol were faster and higher in HeLa, likely due to increased demand for Gln in cancer cells. In contrast, Gln levels in mitochondria were insignificantly different in two cell lines compared to those in cytosol (**Figure 4G**). Interestingly, Gln concentration in mitochondria was generally kept higher than that in cytosol of both cells. Taken together, Q-SHINE provides a simple and effective method of monitoring subcellular Gln levels. Further research utilizing the Q-SHINE system to address the underlying mechanisms how cells distinctly maintain subcellular Gln concentration should provide deeper insights into cellular Gln metabolism.

## 4. Conclusions

In this study, we developed an atypical Gln sensor system by split design of a QBP protein. Q-SHINE exhibits a strong signal-to-noise ratio and a detection range ideally suited to physiological Gln concentrations. The ligand-dependent dimerization of Q-SHINE enables direct measurement of Gln in bio-fluids with comparable sensitivity and higher specificity than a conventional Gln assay kit. Moreover, the performance of genetically encodable Q-SHINE for *in situ* Gln monitoring was demonstrated by comparing cytoplasmic and mitochondrial Gln changes in two distinct cell lines. Indeed, utilizing genetically encoded biosensors to monitor intracellular production or depletion of essential nutrients and metabolites has provided crucial insights by enabling non-invasive diagnosis [33], *in situ* detection of toxins [34], and development of biopharmaceuticals [22]. Collectively, our Gln sensor would facilitate in-depth studies on Gln metabolism and relevant diseases.

## Abbreviations

QBP: glutamine-binding protein
LID: ligand-induced hetero-dimerization
PBP: periplasmic binding protein
BiFC: bimolecular fluorescence complementation
CBP: cysteine-binding protein
HBP: histidine-binding protein

## Acknowledgments

We would like to thank Dr. Sungdo Ha for his valuable support in starting the project. We also thank Jisung Oh and Yebin Won for their help in the experiments.

## Appendix A. Supplementary data

Supplementary data to this article can be found online.

