## Supplementary Materials for "Q-SHINE: a versatile sensor for glutamine measurement via ligand-induced dimerization"

- Supplementary Table 1
- Supplementary Table 2
- Supplementary Figure 1
- Supplementary Figure 2
- Supplementary Figure 3
- Supplementary Figure 4
- Supplementary Figure 5
- Supplementary Figure 6

**Supplementary Table 1.** The protein sequences used throughout the study.

| PBP | Constructs | Amino acid sequence |
|---|---|---|
| QBP | **Split QBP_Lg – mCherry_C** | MHHHHHHLVVATDTAFVPFEFKQGDLYVGFDVDLWAAIAKELKLDYELKPMDFSGIIPALQTKNVDLALAGITITDERKKAIDFSDGYY**TGNG**QQYGIAFPKGSDELRDKVNGALKTLRENGTYNEIYKKWFGTEA**GGGSG**GALKGEIKQRLKLKDGGHYDAEVKTTYKAKKPVQLPGAYNVNIKLDITSHNEDYTIVEQYERAEGRHSTGGMDELYK |
| | **mCherry_N – Split QBP_Sm** | MVSKGEEDNMAIIKEFMRFKVHMEGSVNGHEFEIEGEGEGRPYEGTQTAKLKVTKGGPLPFAWDILSPQFMYGSKAYVKHPADIPDYLKLSFPEGFKWERVMNFEDGGVVTVTQDSSLQDGEFIYKVKLRGTNFPSDGPVMQKKTMGWEASSERMYPED**GGGSG**GLLVMVKANNNDVKSVKDLDGKVVAVKSGTGSVDYAKANIKTKDLRQFPNIDNAYMELGTNRADAVLHDTPNILYFIKTAGNGQFKAVGDSLEALEHHHHHH |
| | **Split QBP_Lg – SmBiT** | MHHHHHHLVVATDTAFVPFEFKQGDLYVGFDVDLWAAIAKELKLDYELKPMDFSGIIPALQTKNVDLALAGITITDERKKAIDFSDGYYTGNGQQYGIAFPKGSDELRDKVNGALKTLRENGTYNEIYKKWFGTEAGSSGGGGSGGGGSSGMVTGYRLFEEIL |
| | **LgBiT – Split QBP_Sm** | MVFTLEDFVGDWEQTAAYNLDQVLEQGGVSSLLQNLAVSVTPIQRIVRSGENALKIDIHVIIPYEGLSADQMAQIEEVFKVVYPVDDHHFKVILPYGTLVIDGVTPNMLNYFGRPYEGIAVFDGKKITVTGTLWNGNKIIDERLITPDGSMLFRVTINSGSSGGGGSGGGGSSGGGLLVMVKANNNDVKSVKDLDGKVVAVKSGTGSVDYAKANIKTKDLRQFPNIDNAYMELGTNRADAVLHDTPNILYFIKTAGNGQFKAVGDSLEAALEHHHHHH |
| | **Q-SHINE_FL** | MSKGEELFTGVVPILVELDGDVNGHKFSVRGEGEGDATNGKLTLKFICTTGKLPVPWPTLVTTLTYGVQCFSRYPDHMKQHDFFKSAMPEGYVQERTISFKDDGTYKTRAEVKFEGDTLVNRIELKGIDFKEDGNILGHKLEYNFNSHNVYITADKQKNGIKANFKIRHNVEDGSVQLADHYQQNTPIGDGPVLLPDNHYLSTQSKLSKDPNEKRDHMVLLEFVTAAGITHGMDELYKSTNSADITSLLVVATDTAFVPFEFKQGDLYVGFDVDLWAAIAKELKLDYELKPMDFSGIIPALQTKNVDLALAGITITDERKKAIDFSDGYYTGNGQQYGIAFPKGSDELRDKVNGALKTLRENGTYNEIYKKWFGTEAGGGSGGALKGEIKQRLKLKDGGHYDAEVKTTYKAKKPVQLPGAYNVNIKLDITSHNEDYTIVEQYERAEGRHSTGGMDELYKGSGEGRGSLLTCGDVEENPGPVSKGEEDNMAIIKEFMRFKVHMEGSVNGHEFEIEGEGEGRPYEGTQTAKLKVTKGGPLPFAWDILSPQFMYGSKAYVKHPADIPDYLKLSFPEGFKWERVMNFEDGGVVTVTQDSSLQDGEFIYKVKLRGTNFPSDGPVMQKKTMGWEASSERMYPEDGGGSGGLLVMVKANNNDVKSVKDLDGKVVAVKSGTGSVDYAKANIKTKDLRQFPNIDNAYMELGTNRADAVLHDTPNILYFIKTAGNGQFKAVGDSLE |
| | **Q-SHINE_FL_ ΔSm** | MSKGEELFTGVVPILVELDGDVNGHKFSVRGEGEGDATNGKLTLKFICTTGKLPVPWPTLVTTLTYGVQCFSRYPDHMKQHDFFKSAMPEGYVQERTISFKDDGTYKTRAEVKFEGDTLVNRIELKGIDFKEDGNILGHKLEYNFNSHNVYITADKQKNGIKANFKIRHNVEDGSVQLADHYQQNTPIGDGPVLLPDNHYLSTQSKLSKDPNEKRDHMVLLEFVTAAGITHGMDELYKSTNSADITSLLVVATDTAFVPFEFKQGDLYVGFDVDLWAAIAKELKLDYELKPMDFSGIIPALQTKNVDLALAGITITDERKKAIDFSDGYYTGNGQQYGIAFPKGSDELRDKVNGALKTLRENGTYNEIYKKWFGTEAGGGSGGALKGEIKQRLKLKDGGHYDAEVKTTYKAKKPVQLPGAYNVNIKLDITSHNEDYTIVEQYERAEGRHSTGGMDELYKGSGEGRGSLLTCGDVEENPGPVSKGEEDNMAIIKEFMRFKVHMEGSVNGHEFEIEGEGEGRPYEGTQTAKLKVTKGGPLPFAWDILSPQFMYGSKAYVKHPADIPDYLKLSFPEGFKWERVMNFEDGGVVTVTQDSSLQDGEFIYKVKLRGTNFPSDGPVMQKKTMGWEASSERMYPEDGGGSGGLLVM |
| | **Q-SHINE_Luc** | MDYKDDDDKGLEGSRMSVLTPLLLRGLTGSARRLPVPRAKIHSLGDPMSVLTPLLLRGLTGSARRLPVPRAKIHSLGDPLVVATDTAFVPFEFKQGDLYVGFDVDLWAAIAKELKLDYELKPMDFSGIIPALQTKNVDLALAGITITDERKKAIDFSDGYYTGNGQQYGIAFPKGSDELRDKVNGALKTLRENGTYNEIYKKWFGTEAGSSGGGGSGGGGSSGMVTGYRLFEEILGSGEGRGSLLTCGDVEENPGPMSVLTPLLLRGLTGSARRLPVPRAKIHSLGDPMSVLTPLLLRGLTGSARRLPVPRAKIHSLGDPVFTLEDFVGDWEQTAAYNLDQVLEQGGVSSLLQNLAVSVTPIQRIVRSGENALKIDIHVIIPYEGLSADQMAQIEEVFKVVYPVDDHHFKVILPYGTLVIDGVTPNMLNYFGRPYEGIAVFDGKKITVTGTLWNGNKIIDERLITPDGSMLFRVTINSGSSGGGGSGGGGSSGGGLLVMVKANNNDVKSVKDLDGKVVAVKSGTGSVDYAKANIKTKDLRQFPNIDNAYMELGTNRADAVLHDTPNILYFIKTAGNGQFKAVGDSLEGGGGSGGGSGEDAKNIKKGPAPFYPLEDGTAGEQLHKAMKRYALVPGTIAFTDAHIEVDITYAEYFEMSVRLAEAMKRYGLNTNHRIVVCSENSLQFFMPVLGALFIGVAVAPANDIYNERELLNSMGISQPTVVFVSKKGLQKILNVQKKLPIIQKIIIMDSKTDYQGFQSMYTFVTSHLPPGFNEYDFVPESFDRDKTIALIMNSSGSTGLPKGVALPHRTACVRFSHARDPIFGNQIIPDTAILSVVPFHHGFGMFTTLGYLICGFRVVLMYRFEEELFLRSXQDYKIQSALLVPTLFSFFAKSTLIDKYDLSNLHEIASGGAPLSKEVGEAVAKRFHLPGIRQGYGLTETTSAILITPEGDDKPGAVGKVVPFFEAKVVDLDTGKTLGVNQRGELCVRGPMIMSGYVNNPEATNALIDKDGWLHSGDIAYWDEDEHFFIVDRLKSLIKYKGYQVAPAELESILLQHPNIFDAGVAGLPDDDAGELPAAVVVLEHGKTMTEKEIVDYVASQVTTAKKLRGGVVFVDEVPKGLTGKLDARKIREILIKAKKGGKIAVYPYDVPDYAGT |
| | **Q-SHINE_BL** | MDYKDDDDKGLEGSRLVVATDTAFVPFEFKQGDLYVGFDVDLWAAIAKELKLDYELKPMDFSGIIPALQTKNVDLALAGITITDERKKAIDFSDGYYTGNGQQYGIAFPKGSDELRDKVNGALKTLRENGTYNEIYKKWFGTEAGGGSGVTGYRLFEEILGSGEGRGSLLTCGDVEENPGPVFTLEDFVGDWEQTAAYNLDQVLEQGGVSSLLQNLAVSVTPIQRIVRSGENALKIDIHVIIPYEGLSADQMAQIEEVFKVVYPVDDHHFKVILPYGTLVIDGVTPNMLNYFGRPYEGIAVFDGKKITVTGTLWNGNKIIDERLITPDGSMLFRVTINSGSSGGGGSGGGGSSGGGLLVMVKANNNDVKSVKDLDGKVVAVKSGTGSVDYAKANIKTKDLRQFPNIDNAYMELGTNRADAVLHDTPNILYFIKTAGNGQFKAVGDSLEGGGSGEDAKNIKKGPAPFYPLEDGTAGEQLHKAMKRYALVPGTIAFTDAHIEVDITYAEYFEMSVRLAEAMKRYGLNTNHRIVVCSENSLQFFMPVLGALFIGVAVAPANDIYNERELLNSMGISQPTVVFVSKKGLQKILNVQKKLPIIQKIIIMDSKTDYQGFQSMYTFVTSHLPPGFNEYDFVPESFDRDKTIALIMNSSGSTGLPKGVALPHRTACVRFSHARDPIFGNQIIPDTAILSVVPFHH |

| | | |
|---|---|---|
| | | GFGMFTTLGYLICGFRVVLMYRFEEELFLRSXQDYKIQSALLVPTLFSFFAKSTLIDKYDLSNL<br>HEIASGGAPLSKEVGEAVAKRFHLPGIRQGYGLTETTSAILITPEGDDKPGAVGKVVPFFEAKV<br>VDLDTGKTLGVNQRGELCVRGPMIMSGYVNNPEATNALIDKDGWLHSGDIAYWDEDEHFFIV<br>DRLKSLIKYKGYQVAPAELESILLQHPNIFDAGVAGLPDDDAGELPAAVVVLEHGKTMTEKEIV<br>DYVASQVTTAKKLRGGVVFVDEVPKGLTGKLDARKIREILIKAKKGGKIAVYPYDVPDYAGT |
| | **mit-Q-SHINE_BL** | MDYKDDDDKGLEGSRMSVLTPLLLRGLTGSARRLPVPRAKIHSLGDPMSVLTPLLLRGLTGS<br>ARRLPVPRAKIHSLGDPLVVATDTAFVPFEFKQGDLYVGFDVDLWAAIAKELKLDYELKPMDF<br>SGIIPALQTKNVDLALAGITITDERKKAIDFSDGYYTGNGQQYGIAFPKGSDELRDKVNGALKT<br>LRENGTYNEIYKKWFGTEAGSSGGGGSGGGGSSGMVTGYRLFEEILGSGEGRGSLLTCGD<br>VEENPGPMSVLTPLLLRGLTGSARRLPVPRAKIHSLGDPMSVLTPLLLRGLTGSARRLPVPRA<br>KIHSLGDPVFTLEDFVGDWEQTAAYNLDQVLEQGGVSSLLQNLAVSVTPIQRIVRSGENALKI<br>DIHVIIPYEGLSADQMAQIEEVFKVVYPVDDHHFKVILPYGTLVIDGVTPNMLNYFGRPYEGIAV<br>FDGKKITVTGTLWNGNKIIDERLITPDGSMLFRVTINSGSSGGGGSGGGGSSGGGLLVMVKA<br>NNNDVKSVKDLDGKVVAVKSGTGSVDYAKANIKTKDLRQFPNIDNAYMELGTNRADAVLHDT<br>PNILYFIKTAGNGQFKAVGDSLEGGGGSGGGSGEDAKNIKKGPAPFYPLEDGTAGEQLHKA<br>MKRYALVPGTIAFTDAHIEVDITYAEYFEMSVRLAEAMKRYGLNTNHRIVVCSENSLQFFMPV<br>LGALFIGVAVAPANDIYNERELLNSMGISQPTVVFVSKKGLQKILNVQKKLPIIQKIIIMDSKTDY<br>QGFQSMYTFVTSHLPPGFNEYDFVPESFDRDKTIALIMNSSGSTGLPKGVALPHRTACVRFS<br>HARDPIFGNQIIPDTAILSVVPFHHGFGMFTTLGYLICGFRVVLMYRFEEELFLRSXQDYKIQSA<br>LLVPTLFSFFAKSTLIDKYDLSNLHEIASGGAPLSKEVGEAVAKRFHLPGIRQGYGLTETTSAILI<br>TPEGDDKPGAVGKVVPFFEAKVVDLDTGKTLGVNQRGELCVRGPMIMSGYVNNPEATNALI<br>DKDGWLHSGDIAYWDEDEHFFIVDRLKSLIKYKGYQVAPAELESILLQHPNIFDAGVAGLPDD<br>DAGELPAAVVVLEHGKTMTEKEIVDYVASQVTTAKKLRGGVVFVDEVPKGLTGKLDARKIREI<br>LIKAKKGGKIAVYPYDVPDYAGT |
| **CBP** | **CBP_Lg-mCherry_C-L3** | MHHHHHHSDSKTLNSLDKIKQNGVVRIGVFGDKPPFGYVDEKGNNQGYDIALAKRIAKELFG<br>DENKVQFVLVEAANRVEFLKSNKVDIILANFTQTPQRAEQVDFSGGLVKKGDKELKEFIDNLII<br>KLGQEQFFHKAYDETLKAHFGDDVKADDVVIEGSSGGGGSGGGALKGEIKQRLKLKDGGHY<br>DAEVKTTYKAKKPVQLPGAYNVNIKLDITSHNEDYTIVEQYERAEGRHSTGGMDELYK |
| | **CBP_Lg-mCherry_C-L5** | MHHHHHHSDSKTLNSLDKIKQNGVVRIGVFGDKPPFGYVDEKGNNQGYDIALAKRIAKELFG<br>DENKVQFVLVEAANRVEFLKSNKVDIILANFTQTPQRAEQVDFTKGGALVKKGDKELKEFIDN<br>LIIKLGQEQFFHKAYDETLKAHFGDDVKADDVVIEGSSGGGGSGGGALKGEIKQRLKLKDGG<br>HYDAEVKTTYKAKKPVQLPGAYNVNIKLDITSHNEDYTIVEQYERAEGRHSTGGMDELYK |
| | **CBP_Sm-mCherry_N** | MKVALGVAVPKDSNITSVEDLKDKTLLLNKGTTADAYFTQNYPNIKTLKYDQNTETFAALMDK<br>RGDALSHDNTLLFAWVKDHPDFKMGIKELGNKDGSSGGGGSGGVSKGEEDNMAIIKEFMRF<br>KVHMEGSVNGHEFEIEGEGEGRPYEGTQTAKLKVTKGGPLPFAWDILSPQFMYGSKAYVKH<br>PADIPDYLKLSFPEGFKWERVMNFEDGGVVTVTQDSSLQDGEFIYKVKLRGTNFPSDGPVM<br>QKKTMGWEASSERMYPEDHHHHHH |
| **HBP** | **mCherry_C-HBP_Lg-L2** | MHHHHHHALKGEIKQRLKLKDGGHYDAEVKTTYKAKKPVQLPGAYNVNIKLDITSHNEDYTIV<br>EQYERAEGRHSTGGMDELYKGGGSGGAIPQKIRIGTDPTYAPFESKNAQGELVGFDIDLAKE<br>LCKRINTQCTFVENPLDALIPSLKAKKIDAIMSSLSITEKRQQEIAFTDKLYKGVNTGMGLRKED<br>NELREALNKAFAEMRADGTYEKLAKKYFDFDVYGG |
| | **mCherry_C-HBP_Lg-L4** | MHHHHHHALKGEIKQRLKLKDGGHYDAEVKTTYKAKKPVQLPGAYNVNIKLDITSHNEDYTIV<br>EQYERAEGRHSTGGMDELYKGGGSGGAIPQKIRIGTDPTYAPFESKNAQGELVGFDIDLAKE<br>LCKRINTQCTFVENPLDALIPSLKAKKIDAIMSSLSITEKRQQEIAFTDKLYADHLGGTGMGLRK<br>EDNELREALNKAFAEMRADGTYEKLAKKYFDFDVYGG |
| | **HBP_Sm-mCHerry_N** | MDSRLVVAKNSDIQPTVASLKGKRVGVLQGTTQETFGNEHWAPKGIEIVSYQGQDNIYSDLT<br>AGRIDAAFQDEVAASEGFLKQPVGKDYKFGGPAVKDEKGSSGGGGSGGVSKGEEDNMAIIK<br>EFMRFKVHMEGSVNGHEFEIEGEGEGRPYEGTQTAKLKVTKGGPLPFAWDILSPQFMYGSK<br>AYVKHPADIPDYLKLSFPEGFKWERVMNFEDGGVVTVTQDSSLQDGEFIYKVKLRGTNFPSD<br>GPVMQKKTMGWEASSERMYPEDHHHHHH |

**Supplementary Table 2.** Comparison of binding energy between the ligands and each domain of PBPs.

| **PBP** | **PDB ID** | **AA** | **ΔΔG [a] (Lg+Sm)** | **ΔΔG (Lg)** | **ΔΔG (Sm)** | **Binding ratio to Lg [b]** | **Binding ratio to Sm** | **Max/Min [c]** |
|---|---|---|---|---|---|---|---|---|
| QBP | 1WDN | Gln | -28.457 | -13.481 | -14.975 | 0.474 | 0.526 | 1.110 |
| CBP | 1XT8 | Cys | -12.410 | -4.911 | -7.499 | 0.396 | 0.604 | 1.525 |
| HBP | 1HSL | His | -11.788 | -7.711 | -4.077 | 0.654 | 0.346 | 1.890 |

[a] The ligand binding energy (ΔΔG, Rosetta Energy Unit) is calculated by RosettaScript ddG mover after energy minimization by REF2015 score function.

[b] Binding ratio is defined as binding energy ratio of ligand for a particular domain. For example, binding ratio to Lg is calculated as ΔΔG (Lg) / ΔΔG (Lg+Sm).

[c] Max/Min is the ratio of maximum to minimum binding ratio value, which represents the extent of balance of ligand-binding by each split domain.

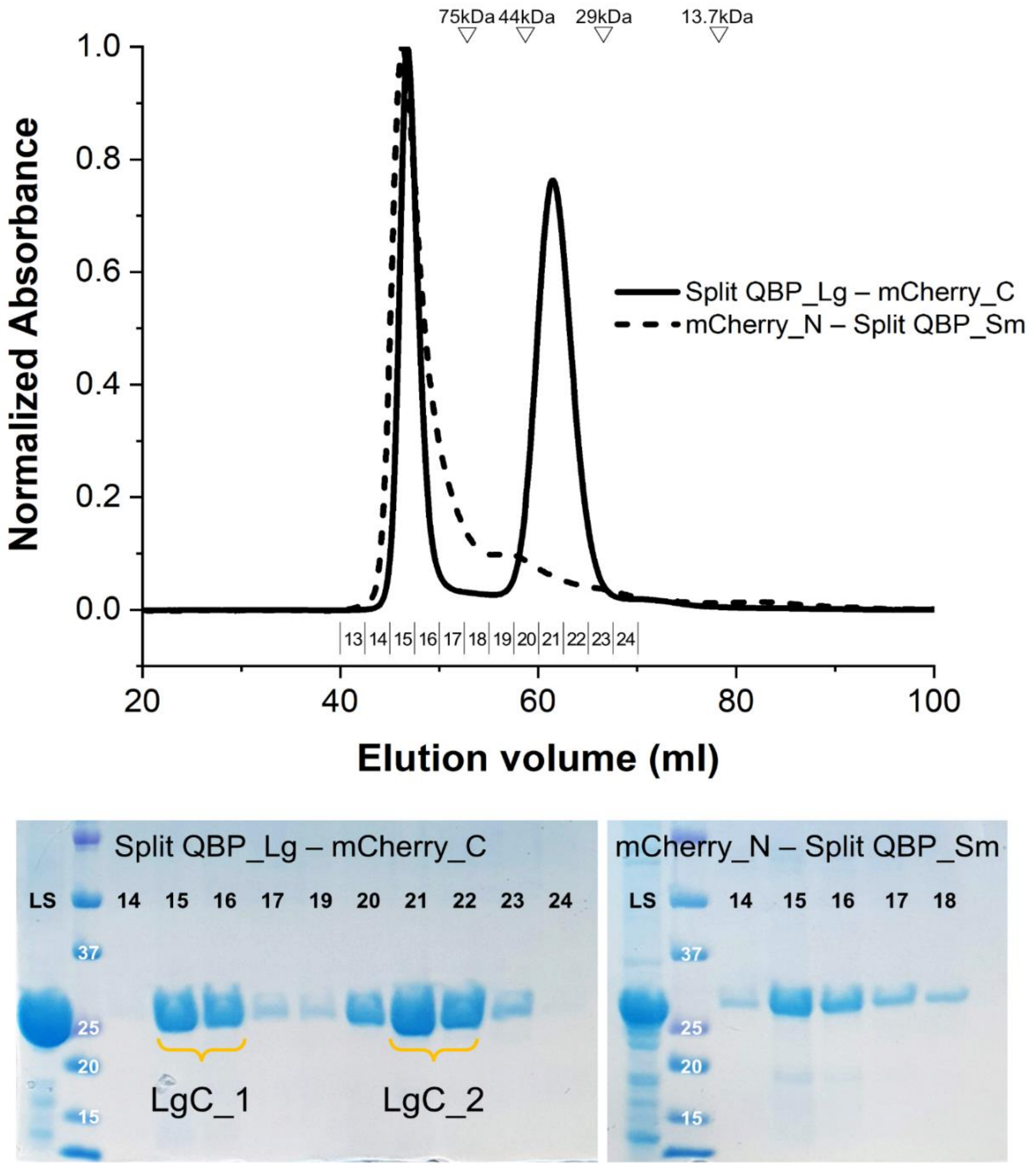


**Supplementary Figure 1.** Analytical gel filtration chromatography profiles of Q-SHINE_Red domains: Split QBP_Lg – mCherry_C (LgC, solid line) and mCherry_N – Split QBP_Sm (SmN, dash line). The original protein (LS) purified from Ni column, and eluted fractions from GPC column (marked from 14 to 24) loaded onto 12% SDS-PAGE. Compared to standard proteins for calibration, the molecular weight of eluted domains was different from the expected value. (24.4 kDa and 29.5 kDa for LgC and SmN, respectively). The two separate peaks (LgC_1(fraction 15/16) and LgC_2(21/22)) of LgC were compared for the measurement.

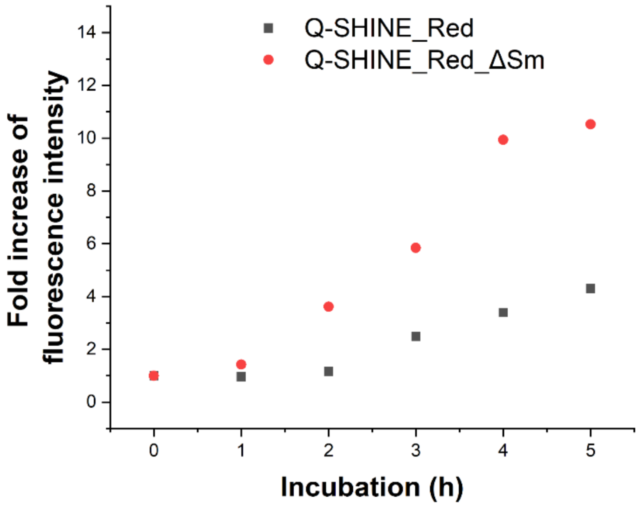


**Supplementary Figure 2.** Comparison of self-assembly propensity of Q-SHINE_Red and Q-SHINE_Red_ΔSm in the absence of glutamine.

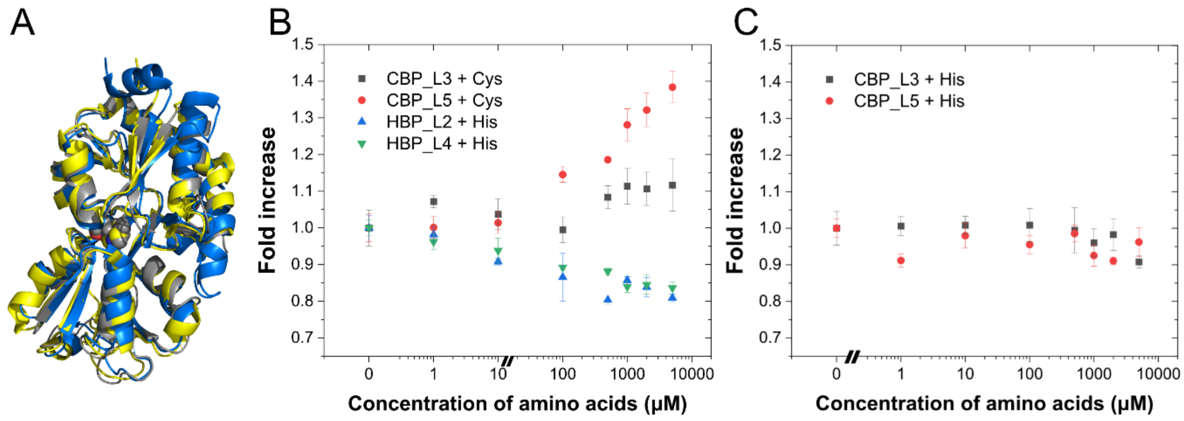


**Supplementary Figure 3.** Split design of PBPs for Cys and His detection. (A) Superposition of QBP (PDB ID: 1WDN; grey), CBP (PDB ID:1XT8, blue) and HBP (PDB ID: 1HSL; yellow). CBP and HBP shows RMSD 1.239 Å and 1.000 Å, respectively. (B) Ligand-dependent BiFC signal from 2 pairs of CBP and HBP were measured with the corresponding concentration of marked amino acids. Results are mean ± s.d. (n = 3) of fluorescence intensity normalized to 0 μM conditions. CBP_L3 and CBP_L5 showed Cys-dependent fluorescence despite their limited dynamic range of the signal compared to that of Q-SHINE_Red. Difference of reactivity between CBP_L3 and CBP_L5 indicates that the design of an optimized linker for noncontinuous split domain is crucial in constituting the binding interface with the ligand. However, His-dependent fluorescence was not detected in both HBP_L2 and HBP_L4. Rather, there was reverse reactivity to the ligand in split HBPs. (C) The split CBPs were incubated with His to test if high concentrations of His affected the self-complementation of split mCherry fragments. There was a trivial effect compared to split HBPs. Thus, the binding of His to the large domain of HBP is supposed to hinder the complementation of two fragments

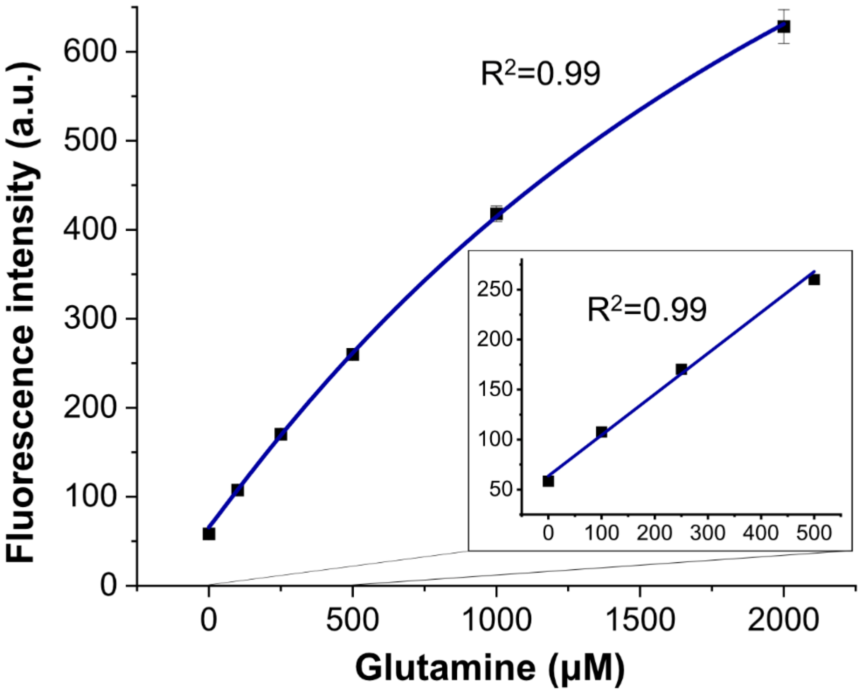


**Supplementary Figure 4.** Standard curve of glutamine solution by Q-SHINE_Red. Although linearity in signal was observed below 500 μM glutamine, we used non-linear standard fitting considering the fact that physiological glutamine concentration normally ranges from 200 to 1400 μM.

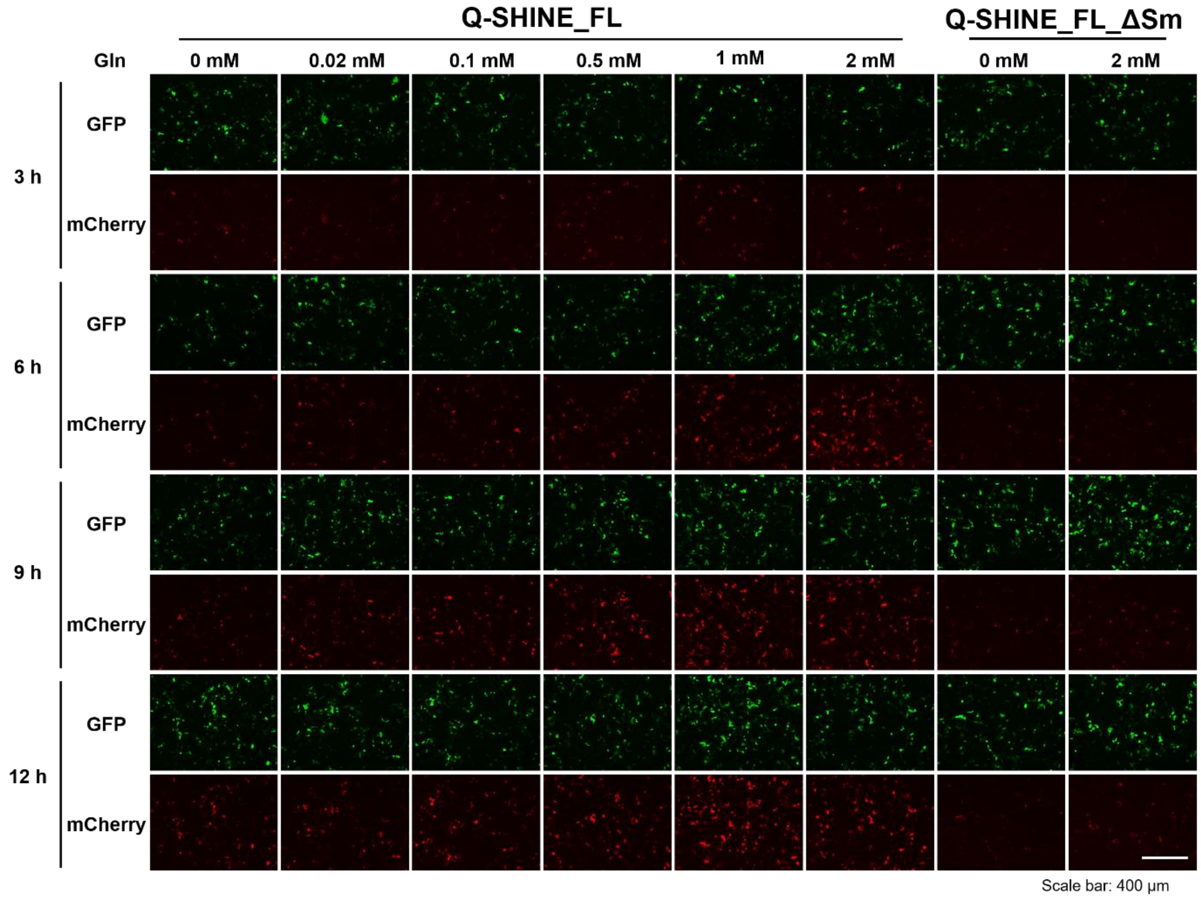


**Supplementary Figure 4.** Separate fluorescence images from green and red channel. Clear difference of red fluorescence is observed from Q-SHINE_FL (A) compared to Q-SHINE_FL_ΔSm (B).

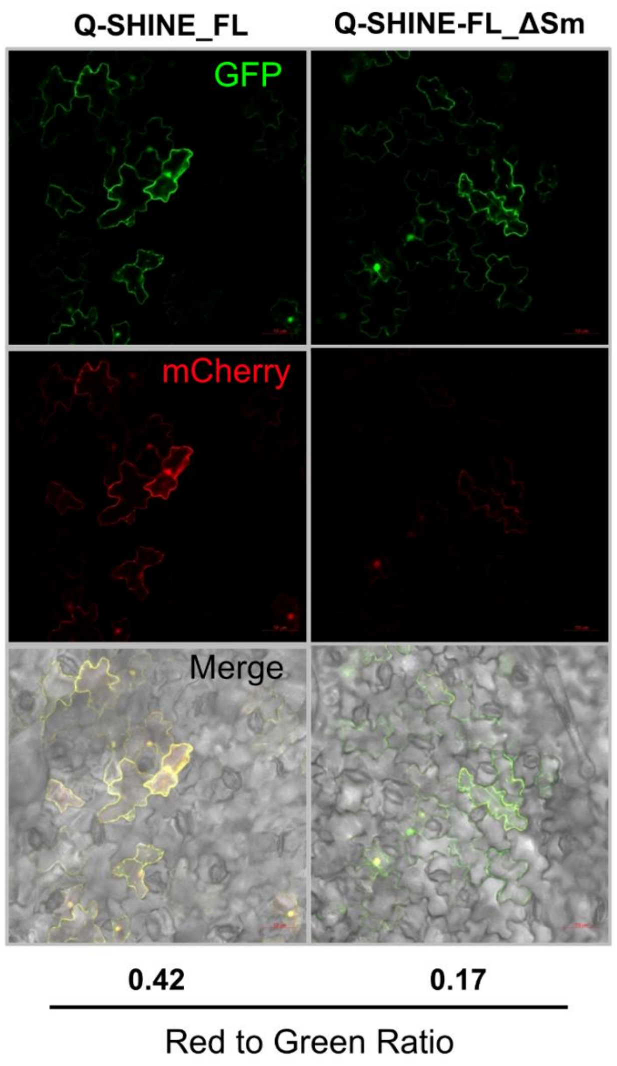


**Supplementary Figure 5**. Detection of Gln in tobacco leaves with genetically encoded Q-SHINE_FL. Red-to-green intensity ratio is marked. Mean of six images; three biological replicates with two technical replicates. Since the endogenous Gln concentration in plant species, including tobacco, is within the range 2.5–20 mM,[33] which is excessive for measurement purposes, we compared fluorescence from Q-SHINE-FL and Q-SHINE-FL_ΔSm.
